# EGR3 Deletion Rescues Developmental and Epileptic Encephalopathy in *Kcna1*-null Mice

**DOI:** 10.1101/2025.06.07.658422

**Authors:** Arindam Ghosh Mazumder, Saifina Karedia, Nandani Adhyapak, Catharina Schimer, John Samuel Bass, Jessica L Kamen, Miranda J Jankovic, Qinglong Miao, Amelia L Gallitano, Alexander B Saltzman, Antrix Jain, Anna Malovannaya, Edward Glasscock, Isamu Aiba, Jeffrey L Noebels, Vaishnav Krishnan

**Affiliations:** Department of Neurology, Baylor College of Medicine, Houston, TX; Department of Biological Sciences, Southern Methodist University, Dallas, TX; Department of Basic Medical Sciences, University of Arizona, Phoenix AZ; Mass Spectrometry Proteomics Core, Advanced Technology Cores, Baylor College of Medicine, Houston TX; Department of Biochemistry and Molecular Pharmacology, Baylor College of Medicine, Houston TX

**Keywords:** Epilepsy, Developmental and epileptic encephalopathy, Seizures, EGR3, KCNA1, Home-cage behavior

## Abstract

*KCNA1* encodes the α-subunit of the voltage-gated potassium channel K_V_1.1. Mutations in K_V_1.1’s pore domain result in developmental and epileptic encephalopathy (DEE), where early life seizures and a culprit lesion synergistically disrupt neurodevelopmental trajectories, resulting in intellectual disability that often presents with disturbances in sleep, sociability and sensory processing. Abnormalities in the subcellular localization of Kv1.1, via mutations in/autoantibodies against LGI1 and CNTNAP2, also give rise to syndromes of epilepsy and neuropsychiatric impairment. Mice with deletions of *Kcna1*^(“-/-“)^ display spontaneous seizures at 2-3 weeks of age and premature mortality. In this study, we applied instrumented home-cage monitoring to examine how aberrations in KCNA1 expression may result in pervasive alterations in spontaneous behavior. Compared to wildtype, *Kcna1*^-/-^ mice displayed a robust multifaceted behavioral syndrome featuring marked nocturnal hyperactivity, insomnia, reduced sheltering, fragmented feeding/drinking rhythms, sensory over-responsivity and diminished wheel-running. In identical recordings, *Kcna1*^+/−^ mice only displayed *increased* sheltering, *Lgi1*^+/−^ mice displayed mild insomnia and *Cntnap2*^-/-^ mice showed home-cage *hypoactivity*. *Kcna1* loss in parvalbumin-positive interneurons (Pv-Cre) resulted in a subtle phenocopy, with mild insomnia accompanied by reduced sheltering behavior, while similar deletions in forebrain pyramidal neurons (Emx1-Cre) or dopaminergic neurons (DAT-Cre) were asymptomatic. Adult-onset conditional deletions of *Kcna1* also produced only mild insomnia 6 weeks later. To survey the molecular landscape in *Kcna1*^-/-^ mice, we conducted a mass spectrometry proteomic analysis of dissected hippocampal tissue (a predominant seizure onset zone and where astrogliosis is observed). This revealed significant upregulations in BDNF (brain-derived neurotrophic factor) and the immediate early transcription factor EGR3 (early growth response-3), which is necessary for the induction of BDNF following electroconvulsive seizures. Heterozygous or homozygous deletions of *Egr3* in *Kcna1*^-/-^ mice resulted in significant survival prolongation, a partial neurobehavioral rescue, and a significant improvement in the frequency of spontaneous seizures *and* spreading depolarization events. These clinical improvements were associated with an amelioration of BDNF induction, hippocampal astrogliosis and proteomic disturbances. Together, these data illustrate how an ion channel that governs excitability at millisecond scales also shapes the spatiotemporal structure of spontaneous behavior at meso- or macroscopic time scales. Our results provide a model and a set of precision endpoints to understand how ictal and interictal features of DEE may be ameliorated by inhibiting the long-term downstream transcriptional alterations imparted by early life seizures.

## Introduction

Developmental and epileptic encephalopathies (DEEs) are characterized by a triad of childhood onset seizures, intellectual disability and neurobehavioral deficits that include disturbances in sleep, communication/social reciprocity, feeding and sensory integration.^1–5^ Cognitive and psychiatric disturbances in DEE are hypothesized to result from two autonomous but intertwined pathophysiological processes, including (i) an inciting lesion (genetic or structural) capable of itself altering neurodevelopmental trajectories, and (ii) periods of frequent early life seizures (ELS), that further aggravate developmental delays.^3,6–11^ In many genetic DEEs, behavioral deterioration may begin prior to seizure onset and may persist despite complete seizure control or spontaneous remission.^2–4,10,12^ The genetic landscape of DEEs^13^ includes pathogenic variants in ion channels (e.g., *Kcnq2, Scn1a, Gabrb3*), transcription factors (e.g., *Arx*), chromatin remodelers (e.g., *Chd2*), cell adhesion molecules (e.g., *Pcdh19*) and kinases (e.g., *Cdkl5*). Preclinical DEE models play a crucial role in causally linking a genetic diagnosis in an affected patient to a syndrome of seizures and neuropsychiatric disturbances, and ultimately designing a clinically informed precision treatment. Mouse models of DEE offer a short lifespan, the opportunity to dissect circuit-specific mechanisms and a modifiable genome that provides unparalleled construct validity for monogenic DEEs. Achieving adequate face validity has been more challenging.^14^ While some models have been shown to faithfully display multiple seizure types^15^, seizures may be difficult to monitor and discern if they arise (and/or result in demise) prior to weaning.^16,17^ In addition, human syndromes are typically associated with heterozygously expressed variants, while mice may only be seizure-prone in the homozygous state^18^, or remain entirely seizure free during the relatively brief durations of laboratory EEG (electroencephalography) observations.^19^

While video-EEG remains the gold-standard technique for clarifying seizure occurrence, attempts to ascertain neurobehavioral phenotypes in these models remain poorly standardized. Separate from tests of cognitive function (e.g., spatial memory, object memory), many laboratories also apply a battery of short behavioral tests to ascertain non-cognitive endpoints (e.g., elevated plus maze [exploratory drive], three-chamber testing [sociability], etc.). The popularity of these testing protocols has recently come under increased scrutiny.^20,21^ First, since they are relatively brief (∼5-60 minutes), these tests provide daytime snapshots of emotionality in an exquisitely nocturnal species. Second, test results are often extrapolated into *human-like* symptoms (e.g., *“anxiety-like”, “schizophrenia-like”*), a practice that stems from their originally intended purpose of achieving rapid drug screening. Third, virtually all these assays require a human experimenter, either to manually score event occurrences (e.g., sniffing, grooming) and/or to manually transfer rodent subjects between their home cages and testing arenas. Effective blinding may be impossible when mutants display obvious phenotypic traits (e.g., microcephaly), and human pheromonal cues can markedly impact behavioral results.^22,23^ Recent advances in videotracking and home-cage instrumentation now permit the collection of more prolonged recordings where multimodal behavioral data is automatically acquired under experimenter-free conditions.^20,24,25^ Using one such commercially available platform, we have previously demonstrated how home-cage monitoring can identify murine-specific features of pervasive neurodevelopment in monogenic epilepsy models^26,27^, following prenatal anticonvulsant exposure^28^, and behavioral trajectories associated with aging.^29^ Home-cage methods allow for repeated longitudinal testing^29,30^ and provide datasets that can be analyzed across multiple time scales.^20^

In this study, we report a syndrome of home-cage behavioral abnormalities in mice with constitutive deletions of *Kcna1* (Kcna1^-/-^). *KCNA1* encodes K_V_1.1, a voltage-gated potassium channel *alpha* subunit that assembles as pore-forming homo- or heterotetramers, together with auxiliary *beta* subunits that modulate channel gating, assembly and trafficking.^31^ Pathogenic *KCNA1* variants cause disorders of paroxysmal hyperexcitability, both in central (episodic ataxia, epilepsy, paroxysmal kinesogenic dyskinesia) and peripheral nervous systems (myokymia).^31^ DEE or DEE-like syndromes have been reported in patients with homozygous or heterozygous loss of function variants in the pore domain.^32–38^ *Kcna1*^-/-^ mice were first described in 1998^39^ and have since been employed by numerous laboratories as a model of severe epilepsy and premature mortality. *Kcna1*^-/-^ mice are nonataxic and reliably display spontaneous seizures beginning at 2-3 weeks of age^39^ with features that suggest a focal onset, beginning with a behavioral pause and head nodding, and progressing to forelimb clonus and tonic-clonic activity^40^. These seizures induce Fos activation within amygdalo-hippocampal circuits^41^, and a proportion of these seizures display a hippocampal onset.^40^ Unmedicated *Kcna1*^-/-^ mice experience high rates of seizure-related sudden death, with few surviving past 8-10 weeks of age.^42,43^ Treatment with a ketogenic diet is only able to delay mortality by a few weeks^44–46^ and many antiseizure medications achieve incomplete seizure control.^47^ In contrast, *Kcna1*^+/−^ mice are seizure-free, breed successfully and have a normal lifespan.^39^ Assessments of behavioral comorbidity in *Kcna1*^-/-^ mice have mainly focused on sleep phenotypes: using EEG-EMG recordings, Kcna1^-/-^ mice exhibit significant *deficits* in both NREM and REM sleep.^48,49^ Here, we reveal that *Kcna1*^-/-^ mice display a set of robust abnormalities in home-cage behavior that co-occur with spontaneous seizures. In contrast, we identify relatively subtle changes in *Kcna1*^+/−^ mice, or in mice with deletions of *Lgi1* or *Cntnap2* (which encode proteins that are necessary for the accurate subcellular localization of K_V_1.1). Using floxed *Kcna1* mice^43^, we achieve only a partial phenocopy of *Kcna1*^-/-^ behaviors following *Kcna1* in parvalbumin-positive interneurons. Through a mass spectrometry proteomic survey of *Kcna1*^-/-^ hippocampi, we identify a significantly altered molecular landscape which features upregulations in a range of neuropeptides and the immediate early transcription factor EGR3 (early growth response 3) an immediate early transcription factor known to be induced by electroconvulsive seizures.^50^ We show that deletions of *Egr3* in *Kcna1*^-/-^ mice achieve ameliorations in neurobehavioral abnormalities, proteomic changes, as well as seizure frequency and premature mortality.

## Methods

### Animals and Treatments

All animal protocols were approved by the Baylor College of Medicine Institutional Animal Care and Use Committee. All data elements as per ARRIVE guidelines are provided. Mice were bred and weaned under standard vivarium conditions with a 12:12h light cycle (1700 lights OFF). Genotyping was conducted at postnatal day 21 using tail DNA samples. Our study includes mice of the following lineages: *Kcna1*^+/−^ (Tac:N:NIHS-BC)^39,45,46,51^, floxed *Kcna1* (*Kcna1^fl^*^/fl^, C57BL/6)^43,52–54^, *Egr3*^+/−^ (C57BL/6)^55–57^, *Lgi1*^+/−^ (C57BL/6, JAX: 030640)^58^ and Cntnap2^+/−^ (C57BL/6, JAX: 017482)^59^. We utilized the following Cre-expressing mice: CAGGCreER^TM^ (JAX: 004682)^60^, DAT-Cre (dopamine transporter, JAX: 006660)^61^, PV-Cre (parvalbumin, JAX: 017320)^62^ and EMX1-Cre (JAX: 005628)^63^. Primer sequences for genotyping are provided in Supplementary Material. *Lgi1*^+/−^ mice and *Cntnap2*^+/−^ mice were bred to achieve *Lgi1*^+/−^ *Cntnap2*^+/−^ mice. Double-heterozygous *Egr3^+/−^Kcna1^+/−^* mice were bred to compare behavioral profiles for *Egr3*^+/+^ *Kcna1*^+/+^, *Egr3*^+/+^ *Kcna1*^-/-^, *Egr3*^+/−^ *Kcna1*^-/-^ and *Egr3*^-/-^ *Kcna1*^-/-^ mice. *Egr3*^-/-^*Kcna1*^+/+^ were also included in a proteomic survey, To examine the consequence of adult-onset *Kcna1* deletion, ∼7-week-old CAGGCre.ER^TM^+.*Kcna1^fl^*^/fl^ and CAGGCreER- *Kcna1^fl^*^/fl^ littermates received five daily intraperitoneal injections of tamoxifen dissolved in corn oil (0.5ml/kg).^64^

### Home-cage monitoring

Recordings of home-cage behavior were conducted as previously described^26,28,29,65^ in 6-8 week old-mice unless otherwise specified. At the start of monitoring, mice were singly housed in Noldus Phenotyper® home-cages (30x30x47cm), fitted with a top unit containing an infrared (IR) camera and IR bulbs), an infrared-lucent shelter, two lickometer-connected water-spouts (providing fresh drinking water with 0% Vs 0.8% sucrose), and a beam-break metered food hopper (Fig. 1A). Home-cages were housed within a temperature- and humidity-controlled satellite facility fitted to a 12:12 light cycle (OFF at 1700). Video-recording and tracking was conducted via Ethovision XT 14. We employed a modular design^26,28,29,65^, abbreviated to at least capture (i) at least two 23h-long “baseline recordings” (from 1400 → 1300), (ii) responses to the “light spot test” (1900-2000)^66^ and (iii) measurements of voluntary wheel-running on the final day (from 1400 → 1300). CAGGCre.ER^TM^ - Kcna1^fl/fl^ and CAGGCre.ER^TM^+. Kcna1^fl/fl^ did not undergo light spot or wheel running evaluations. Mouse centerpoint X-Y coordinates sampled at 15Hz were automatically tallied to measure horizontal displacement (e.g., cm/min, cm/hour), heatmaps of position probability, shelter engagement time and were also applied to estimate the timing and duration of “sleep” bouts during baseline recordings using established behavioral definitions (≥40s of contiguous immobility^67^). From lickometers and feeding meters, we extracted both the duration and occurrences of licking and feeding bouts. Wheel rotations/min were tallied using a magnetic wheel sensor (Noldus).

**Figure 1.**
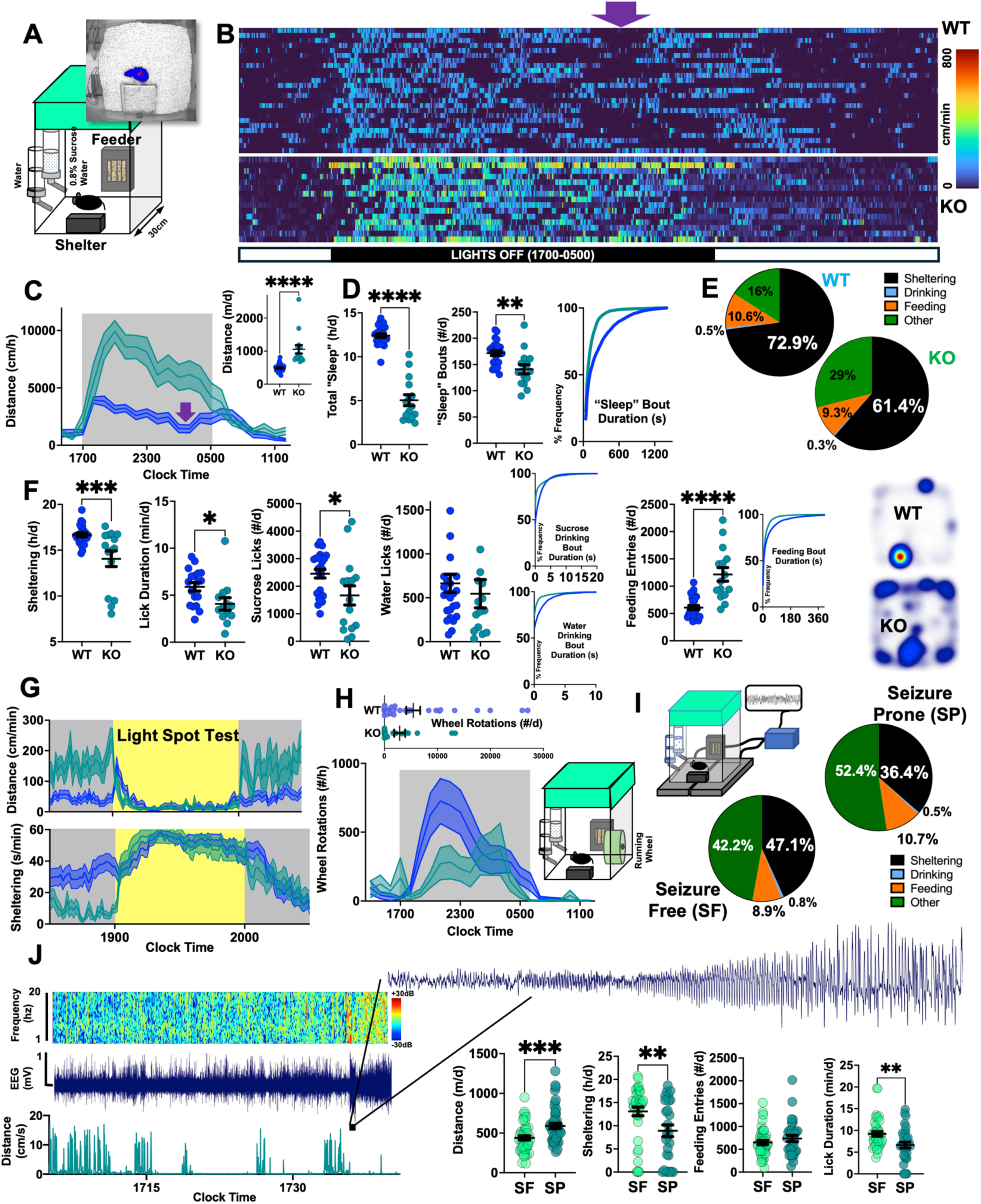
*Kcna1*-/- mice display multiple aberrations in home-cage behavior in concert with spontaneous seizures. (A) Mice are studied within square (30x30cm) home-cages featuring an infrared-(IR)lucent shelter, two lickometered water sources, a food hopper, an aerial IR camera and IR bulbs. (B) Raster plot, with every row depicting a single mouse and where every cell depicts the horizontal distances accumulated within a minute of the day. Purple arrow: a generally synchronized rest period is observed in WT mice (n = 19, 10 female) during the mid-active phase, and which is absent in *Kcna1*-/- mice (n = 13, 7 female). (C) *Kcna1*-/- mice traverse greater distances over baseline recordings (hour x genotype, F21, 798 = 10.89, p<0.0001). (D) This effect is associated with a reduction in behaviorally defined “sleep” (see Methods), fewer “sleep” bouts that were also shorter in duration (KS p<0.0001). (E) Time budgets for sheltering, drinking, feeding and other behaviors. (F) In addition to lower sheltering rates, *Kcna1*-/- mice spend less time licking, accumulate fewer sucrose but not water licks and achieve greater feeding entries. Bout durations for sucrose and water licking, as well as feeding are significantly shorter (left shifted, KS p<0.0001 for all). Representative heat maps of position probability are shown. (G) Responses to an hour-long light spot test applied at 1900. (H) *Kcna1*-/- mice accumulate fewer wheel rotations (F22, 731 = 2.64, p<0.0001). (I) Wireless EEG achieved within the home-cage by combining two baseplate receivers. During seizure prone days (SP), *Kcna1*-/- mice display more pronounced reductions in sheltering compared with seizure-free days. (J) Frequency spectrogram, raw EEG and high-resolution actogram during a representative spontaneous electrographic seizure. On seizure-prone days, *Kcna1*-/- mice accumulated significantly *greater* horizontal distances, lower sheltering rates and lick durations. Mean + s.e.m shown for all. *, **, ***, **** depict p<0.05, <0.01, <0.001 and <0.0001 respectively. KS (Kolmogorov-Smirnov test).

### Data Visualization and Statistical Analysis

We employed sample sizes comparable to previously published results using our home-cage system. Comparisons were always made against littermate controls. When presenting data from “baseline recordings”, the second baseline day is shown. All graphs and statistical comparisons were generated on Prism GraphPad 10, while heatmaps were generated via Matlab. Contrasts between two or among three or more groups were analyzed by unpaired student’s T tests and one-way ANOVA tests, respectively. Post-hoc comparisons for one-way ANOVA were conducted via Tukey’s multiple comparisons test. Significant *genotype x time* interactions with repeated measures (e.g., Fig. 1C) were assessed using a mixed model approximating a 2-way ANOVA.^26–28^ The Kolmogorov-Smirnov (KS) test was employed to compare cumulative frequency distributions of bout lengths (e.g., Fig. 1D). The log-rank (Mantel-Cox) test was employed to compare survival curves. All graphs depict mean ± SEM.

Video-EEG, mass spectrometry proteomics, immunohistochemistry, ELISA and qRT-PCR methods are provided in supplemental material.

## Results

### Kcna1^-/-^ Mice Display Widespread Aberrations in Home-Cage Behavior

When compared against wildtype (WT) littermates within our instrumented home-cages (Fig. 1A) during baseline recordings (see Methods), 5-week-old *Kcna1*^-/-^ mice displayed profound nocturnal hyperactivity (Fig. 1B, C), achieving an approximate doubling in daily horizontal distance (∼1000m vs ∼480m/d) within a 30x30cm square home-cage. WT mice displayed a characteristic^24^ and relatively synchronized dip in activity during the mid-active phase, which was absent in *Kcna1*^-/-^ mice. In conjunction with hyperactivity, *Kcna1*^-/-^ mice displayed significant reductions in “sleep” (enquoted, since sleep here is determined behaviorally), and is consistent with previous reports that have employed beam-break actigraphy assessments^44^ and traditional EEG/EMG.^48,49^ “Sleep” bouts were fewer in number and significantly shorter in duration (Fig. 1D). Time budgets^68^ calculated over the same baseline recordings revealed significant reductions in sheltering and licking durations that could not be explained by increased feeder engagement (Fig. 1E, F). Total sucrose licks (but not water licks) were significantly fewer in *Kcna1*^-/-^ mice and licking bouts across both liquid sources were significantly shorter in duration (Fig. 1F). While *Kcna1*^-/-^ mice displayed a greater number of feeding bouts, bout *durations* of feeding were also significantly shorter (Fig. 1F). Next, to examine how *Kcna1*^-/-^ mice respond to an aversive stimulus applied within the *same* home-cage, we presented an hour-long light spot^28,66^ during the early nocturnal phase. As shown in Fig. 1G, WT mice displayed an initial burst in activity that gradually recedes as shelter engagement becomes more sustained. In contrast, *Kcna1*^-/-^ mice responded with brisk hypoactivity and shelter engagement in response to the stimulus, with an equally brisk resumption of hyperactivity following stimulus cessation. The following day, we deployed a running wheel within this home-cage. *Kcna1*^-/-^ mice accumulated significantly fewer wheel rotations over this trial (Fig. 1H), potentially signaling a hedonic deficit (akin to reduced sucrose drinking). Finally, to broadly examine how inter-daily variations in seizure frequency may be linked to fluctuations in the same home-cage metrics, we deployed wireless EEG implants in a separate cohort of *Kcna1*^-/-^ mice and conducted simultaneous home-cage recordings. On average, seizure-prone days (containing at least one spontaneous seizure) featured significantly *greater* total daily distances and reduced sheltering (Fig. 1I, J). EEG implantation itself imposed a marked reduction in home-cage activity levels (compare with Fig. 1C). Nevertheless, we identified a weak but significant positive correlation between seizure frequency (range 0 −22/d) and total daily distances (r = +0.37, p<0.001). Together, these findings reveal a syndrome of behavioral disturbances that is phenotypically compatible with an encephalopathy, featuring core deficits in arousal and sleep, as well as more diffuse derangements in sensory responsivity, amotivation and fragmented feeding/drinking rhythms. Further, our EEG results suggest that the severity of encephalopathy in *Kcna1*^-/-^ mice maybe dynamically modulated by (or modulates) day to day variations in seizure risk.

### On the Impact of Gene Dosage and Alterations in Subcellular Distribution

Heterozygous loss of function variants in *KCNA1* can result in either episodic ataxia, myokymia and/or epilepsy in humans. *KCNA1* variants within the pore domain are most tightly linked to epilepsy and DEE.^31^. *Kcna1*^+/−^ mice are seizure free^39^, but exhibit a lowered seizure threshold when exposed to fluorothyl^39^, and are also more prone than WT mice to develop electrographic seizures following perinatal hypoxia.^69^ To examine how reductions in *Kcna1* expression may shape home-cage behavior in the absence of spontaneous seizures, we compared a separate cohort of *Kcna1*^+/−^ mice and WT littermates (Fig. 2A, Fig. S1A). *Kcna1*^+/−^ mice exhibited *greater* sheltering times without an increase in total daily “sleep” or overall activity levels. Light spot testing revealed a more sustained sheltering response in Kcna1^+/−^ mice, extending for several minutes beyond stimulus cessation (Fig. 2A). These results suggest that any potential linear relationships between *Kcna1* gene dosage and home-cage behavior abnormalities are either absent or distorted by spontaneous seizures in *Kcna1*^-/-^ mice.

**Figure 2.**
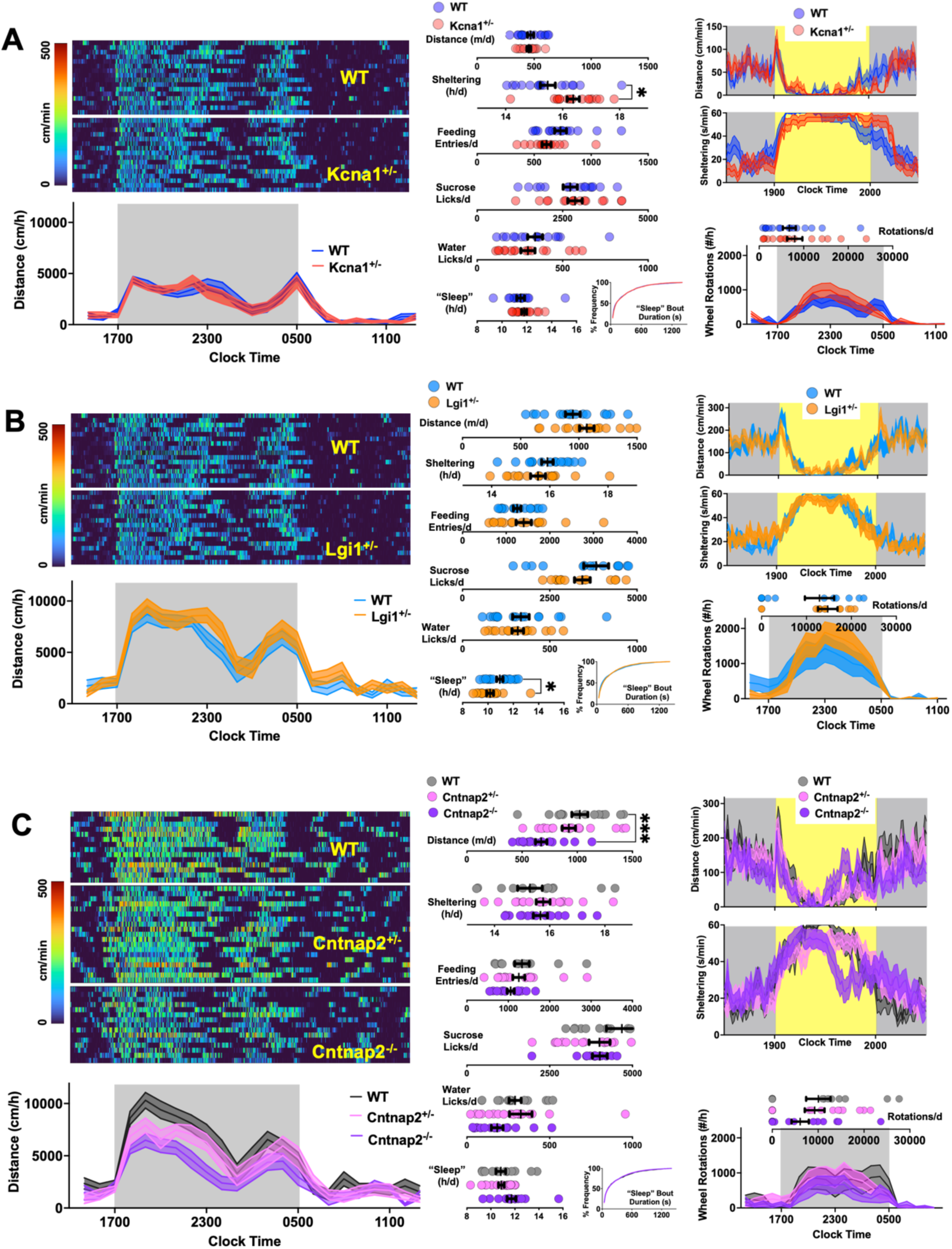
Home-cage behavior in *Kcna1*+/−, *Lgi1*+/−, *Cntnap2*+/− and *Cntnap2*-/- mice. Each panel depicts raster plots of locomotor activity (top left), hourly distances accumulated (bottom left), total metrics for the second baseline recording (center), and averaged light spot and wheel running responses (right). (A) Compared with WT littermates (n=16, 8 female), *Kcna1*+/− mice (n=16, 8 female) display a mild increase in overall sheltering time during baseline recordings. (B) Compared with respective WT littermates (n=16, 8 female), *Lgi1*+/− mice (n=16, 8 female) display a mild but significant reduction in total daily “sleep”. (C) *Cntnap2*-/- mice (n=19, 6 female) feature significant reductions in total daily distances compared with *Cntnap2*+/− (n = 17, 10 female) or littermate WT mice (n=16, 7 female). Mean + s.e.m shown for all. *, *** depict p<0.05 and <0.001 <0.0001 respectively. KS (Kolmogorov-Smirnov).

The accurate subcellular localization of KCNA1 is governed in part by two other proteins that are encoded by epilepsy genes: LGI1 and CASPR2. LGI1 is a secreted protein that modulates neuronal excitability by binding its cognate receptors, ADAM22/ADAM23.^70^ Heterozygous variants in *LGI1* that impair protein secretion or receptor binding give rise to autosomal dominant lateral temporal lobe epilepsy with a penetrance of ∼67%.^71^ Auto-antibodies against LGI1 are among the most frequently identified cause of autoimmune encephalitis^72^, where progressive neurocognitive deterioration is accompanied by dystonic seizures. *Lgi1*^+/−^ mice have a normal lifespan and fertility, while *Lgi1*^-/-^ mice develop spontaneous seizures in the second postnatal week and die by post-natal day 20.^58,73^ Lgi1 loss is thought to contribute to seizures by regulating excitatory synaptic transmission^74,75^ and its role in maintaining the expression of KCNA1 at axon terminals^76^ and the axon initial segment.^77^ Membrane fractions from *Lgi1*^+/−^ mice display ∼50% reduction in KCNA1 expression, and patient-derived anti-LGI1 antibodies effectively lower total and synaptic levels of KCNA1 expression.^78^ When compared against their respective WT littermates within home-cages, Lgi1^+/−^ mice displayed mild insomnia without hyperactivity (Fig. 2B, Fig.S1B), but were otherwise indistinguishable during baseline trials, light spot and wheel running trials.

CASPR2, encoded by *CNTNAP2*, is a trans-synaptic membrane protein that is necessary for the clustering of KCNA1 subunits to the juxtaparanodal region of myelinated axons^79,80^ and for normal patterns of neuronal migration.^59,81,82^ In humans, biallelic loss of function variants of *CNTNAP2* give rise to DEE, featuring autism and focal cortical dysplasia, while the pathogenicity of *Cntnap2* haploinsufficiency remains controversial.^83,84^ Given strong peripheral and central nervous system expression, auto-antibodies against Caspr2 result in autoimmune encephalitis that is frequently accompanied by neuromyotonia and dysautonomia.^72,85^ While *Cntnap2* ^-/-^ mice exhibit normal nerve conduction velocities, they display reductions in social interactions/vocalizations and develop spontaneous seizures beginning at ∼6 months of age.^59,79,81^ Mice with deletions of *Cntnap2* and/or infusions of patient-derived anti-Caspr2 display hypersensitivity to noxious stimuli through the downregulation of K_V_1.1 in membranes of dorsal root ganglion neurons^86^. Compared to WT mice within home-cages, *Cntnap2*^-/-^ mice accumulated significantly *lower* total horizontal displacement during baseline recordings, replicating prior results^87^ (Fig. 2C, Fig. S1C). This effect came with more prolonged drinking and feeding durations (Fig. S1C), but without changes in “sleep” amount and timing. WT, *Cntnap2*^+/−^ and *Cntnap2*^-/-^ mice displayed similar initial responses to light spot testing, while Cntnap2^-/-^ mice featured early shelter exit, suggesting a state of lowered risk aversion. WT and double-heterozygous *Cntnap2*^+/−^*Lgi*^+/−^ mice were also indistinguishable across our home-cage metrics (Fig. S1D). Together, these results demonstrate that *Kcna1* haploinsufficiency or genetic lesions that disturb the subcellular localization of KCNA1 can also perturb metrics of home-cage behavior in directions that are quite distinct from *Kcna1*^-/-^ mice.

### Effects of Temporal and Spatially Precise Deletions of Kcna1

Next, we explored whether widespread deletions of *Kcna1* achieved beyond early developmental timepoints are sufficient to recapitulate the behavioral abnormalities in *Kcna1*^-/-^ mice. We adopted an established strategy that utilizes the expression of tamoxifen-inducible Cre-recombinase under the control of a ubiquitous promoter^64^ crossed with floxed *Kcna1* mice (*Kcna1^fl^*^/fl^)^43,88^. As shown in Fig. 3A, CAGGCre.ER^TM^ positive and negative *Kcna1^fl^*^/fl^ mice displayed similar home-cage metrics prior to tamoxifen exposure. Six weeks later, tamoxifen exposure produced a significant, > 90% reduction in *Kcna1* mRNA levels (Fig. S2A). At this time point, CAGGCre.ER^TM^+ *Kcna1^fl^*^/fl^ mice displayed a mild but significant reduction in total daily “sleep” with subtle reductions in sleep bout durations, but were otherwise indistinguishable from CAGGCre.ER^TM^-negative littermates (Fig. 3A, Fig. S2A). Thus, a deletion of *Kcna1* delayed until adulthood is likely to elicit compensatory mechanisms that maintain behavioral homeostasis.

**Figure 3.**
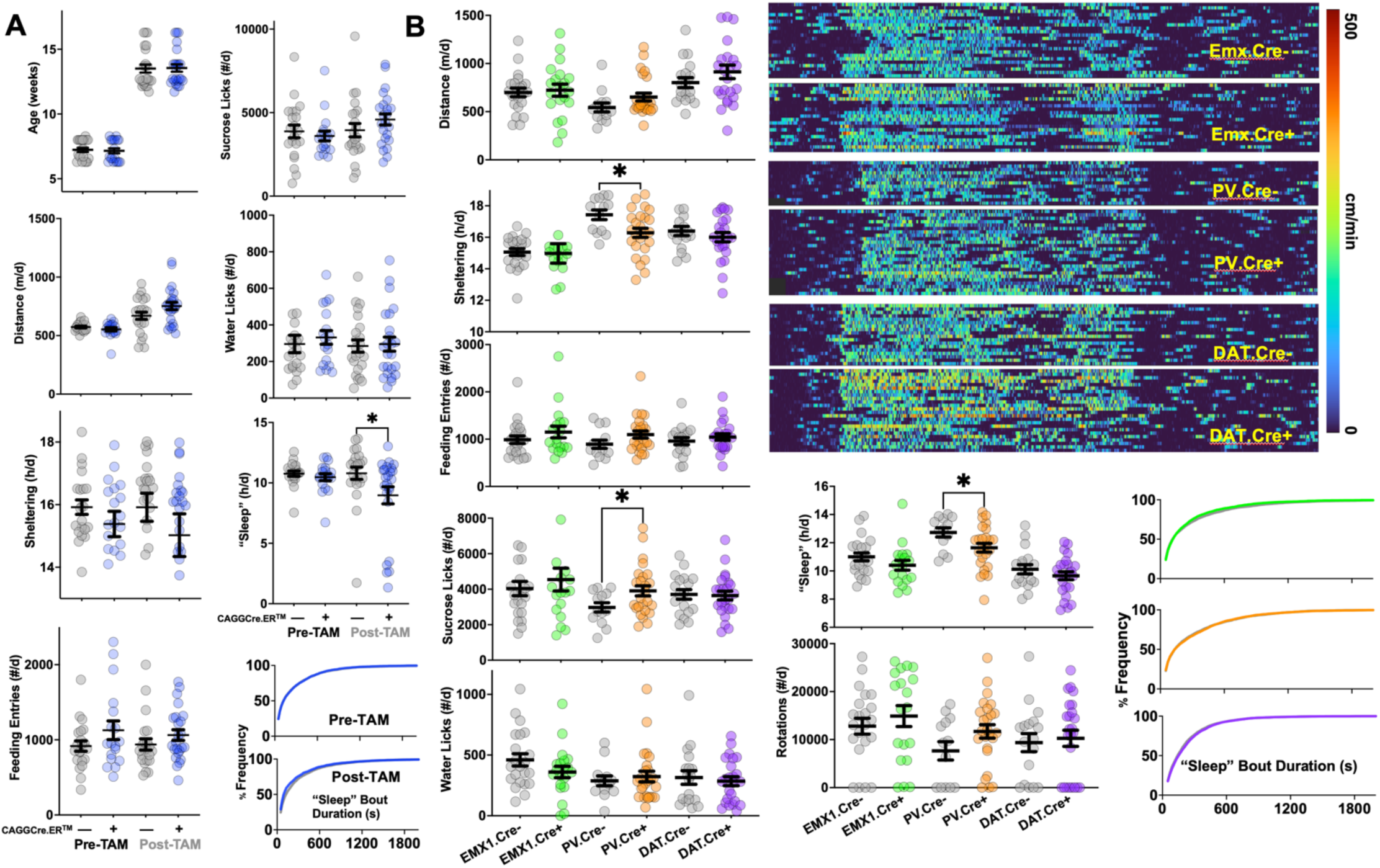
Effects of delayed or cell-type specific genetic deletions of *Kcna1*. (A) Prior to tamoxifen induction (“Pre-TAM”), CAGGCre.ERTM+ *Kcna1fl*/fl (n=19, 9 female) and CAGGCre.ERTM- *Kcna1fl*/fl mice (n=21, 11 female) were indistinguishable on baseline recordings. 6 weeks following tamoxifen exposure, compared with CAGGCre.ERTM- *Kcna1fl*/fl mice (n=23, 10 female), CAGGCre.ERTM+. *Kcna1fl*/fl mice (n=24, 12 female) displayed a mild but significant reduction in total daily “sleep”, as well as a subtle reduction in “sleep” bout durations (KS p<0.05). (B) Selective deletions of *Kcna1* from PV+ interneurons result in *Kcna1*-/- like reductions in sheltering and “sleep”. All remaining home-cage metrics were unchanged across groups. Cre-negative mice (grey) were always compared against respective Cre-positive littermates. Sample sizes: Emx1Cre- (n=22, 6 female), Emx1Cre+ (n=20, 13 female), PVCre- (n=13, 10 female), PVCre+ (n=25, 13 female), DATCre- (n=18, 10 female) and DATCre+ (n=21, 8 female). Mean + s.e.m shown for all. *depicts p<0.05. KS (Kolmogorov Smirnov test)

Conditional deletions of *Kcna1* in excitatory forebrain neurons of the neocortex, hippocampus, amygdala and certain vagal circuits^89^, achieved via the Emx1-Cre driver are sufficient to give rise to spontaneous seizures and premature mortality, albeit with a milder phenotype compared with *Kcna1*^-/-^ mice.^54^ To examine how *Kcna1* loss in specific neuronal populations may phenocopy aspects of the *Kcna1*^-/-^ neurobehavioral syndrome, we recorded the same home-cage metrics in *Kcna1^fl^*^/fl^ mice with or without the concurrent expression of Cre-recombinase in Emx1-expressing, parvalbumin (PV)-expressing or dopamine-transporter (DAT)-expressing cells. KCNA1 plays a key role in controlling the excitability of fast-spiking interneurons.^90–92^ Similarly, Kv1 channels also function to restrain dopamine release from mesencephalic dopaminergic neurons^93,94^ where their expression is necessary for the central stimulatory effects of amphetamine.^95,96^ Emx1- and PV-Cre positive mice displayed a marked reduction in hippocampal *Kcna1* mRNA levels (Fig. S2B, C). Mice with *Kcna1* deletions in Emx1+ or DAT+ neurons, were indistinguishable from their respective Cre-negative littermates. In contrast, mice with PV-specific deletions of Kcna1 displayed reductions in total daily “sleep” and sheltering rates, together with a significant *increase* in sucrose drinking (Fig.3B, Fig. S2B-D). Responses to light spot stimulation or wheel-running were similar in all three conditions (Fig. S2B-D). We confirmed a significant downregulation of hippocampal *Kcna1* mRNA in CAGGCre.ER^TM,^ Emx1Cre and PVCre expressing mice (Fig. S2A-C). Together, these findings suggest that loss of *Kcna1* in excitatory cortical and limbic circuits may serve to selectively enhance seizure risk, while similar deletions of *Kcna1* in PV interneurons may drive alterations in sleep and sheltering behavior in *Kcna1*^-/-^ mice. *Kcna1* deletions in either cell type alone were insufficient to recapitulate the robust hyperactivity and bout fragmentation observed with constitutive deletions of *Kcna1*.

### Changes to the Hippocampal Proteome in Kcna1^-/-^ Mice

Next, to reveal insights into the molecular mechanisms that may directly *mediate* all or parts of the *Kcna1*^-/-^ neurobehavioral syndrome, we conducted microscale label-free proteome profiling of snap-frozen dissected whole hippocampal tissue. After correcting for multiple comparisons, we identified 150 proteins that were significantly up-(60) or down-regulated (90) in expression (Fig. 4A, Supplemental table 1)). Downregulated proteins included KCNA1, as well as several proteins where reduced expression or loss of function has also been linked to DEE, including WDR45^97^, GRIN2A^98^, CASPR2^84^, SYNGAP1^99^, KCND2^100^ and GRIA2.^101^ Among upregulated proteins, we identified a series of neuropeptides that have been linked to temporal lobe epilepsy, including several members of the granin family (SCG2, SCG3, CHGB)^102–104^, preproenkephalin (PENK)^105^, NPY^106^, CCK^107^ and BDNF.^108^ Increases in the hippocampal expression of enkephalin and NPY have been previously reported in the megencephaly (*mceph)* mouse, which has a frameshift deletion that truncates *KCNA1.*^109^ While each of these targets are independently worthy of validation and follow up, we focused on an upregulated transcription factor, EGR3, as a potential direct link between spontaneous seizures and long-term maladaptive transcriptional changes.

**Figure 4.**
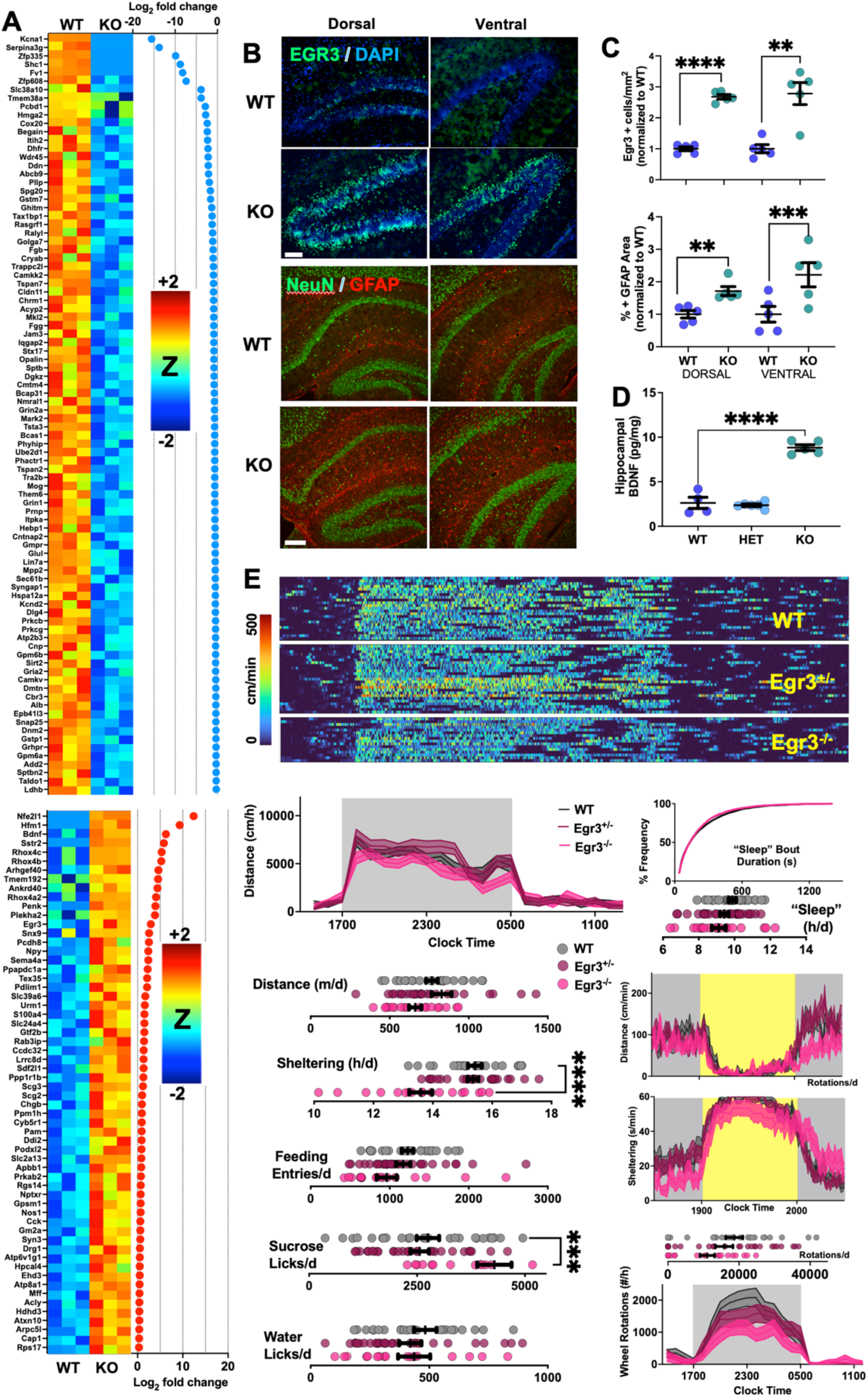
Hippocampal tissues in *Kcna1*-/- mice display an induction of EGR3 and BDNF. (A) Z-normalized intensity-based absolute quantification (iBAQ) scores for proteins that are significantly downregulated (top) and upregulated (bottom) after correcting for false discoveries. N=3 biological replicates are shown for each group. See Supplemental Table 1 for full protein lists. (B) Confirmation by immune- histochemistry of EGR3 and GFAP induction in both ventral and dorsal hippocampal fields (n=5/genotype, scale bar = 50um). (C) Quantification of B. (D) Confirmation of BDNF induction via ELISA (n=5/group). (E) Compared with WT mice (n=25, 13 female) and *Egr3*+/− mice (n=30, 14 female), *Egr3*-/- mice (n=18, 8 female) displayed significantly lower sheltering rate and increased sucrose licks during baseline recordings, without significant increases in total daily “sleep”. Light-spot and wheel- running responses were similar. Mean + s.e.m shown for all. **, ***, **** depict p<0.01, <0.001 and <0.0001 respectively.

EGR3 belongs to a family of immediate early genes (EGR1-4) with three zinc-finger motifs that bind to a canonical EGR DNA response element.^50^ Intronic variants in *EGR3* have been linked to schizophrenia, but have not yet met genome-wide significance.^110–113^ Reductions in EGR3 expression have also been observed in postmortem samples of prefrontal cortex and hippocampus from patients with schizophrenia.^114^ Following electroconvulsive seizures (ECS), EGR3 expression is induced in hippocampal dentate granule cells^115,116^ and is necessary for ECS-induced increases in BDNF expression^56^ and dendritic remodeling.^117^ *Egr3*^-/-^ mice display a mild gait ataxia due to muscle spindle agenesis^57^ and other behavioral phenotypes that have been extrapolated as *“schizophrenia-like”*: heightened aggression^118^, open-field hyperactivity^119^, exaggerated startle responses^119^, mild insomnia^55^ and memory impairments that correlate with deficits in both hippocampal long term depression *and* potentiation.^119,120^ Based on these observations, we hypothesized early onset seizures in *Kcna1*^-/-^ mice result in a vicious cycle of inductions in EGR3 and the pro-epileptic neurotrophin BDNF^108,121,122^ that could sustain seizure risk and EGR3 levels. If EGR3 hypofunction is capable of broadly disrupting neurocognitive function, we reasoned that pathological *increases* in EGR3 activity may also exert deleterious effects on behavior, as shown in models of *Mecp2* ^123^ or *Kctd13*^124^ mutations.

We validated the induction of EGR3 on a separate cohort of WT and Kcna1^-/-^ mice using immunohistochemistry, confirming a ∼3-fold increase in EGR3 positive cells in dentate granule cells in both dorsal (anterior) and ventral (posterior) hippocampal regions (Fig. 4B, C). GFAP staining, as a marker of astrogliosis^51^, was also elevated (Fig. 4B, C) and was also identified in our unbiased proteomic screen (supplemental table 1). Using ELISA in separate mice, we confirmed that whole hippocampal lysates *Kcna1*^-/-^ mice contained significantly increased levels of BDNF (Fig. 4D). In recordings of home-cage behavior (Fig. 4E, Fig. S3A), *Egr3*^+/−^ and *Egr3*^-/-^ mice exhibited levels of activity and “sleep” during baseline recordings that were comparable to WT littermates. *Egr3*^-/-^ mice spent a significantly lower time in the shelter and a significantly elevated number of sucrose licks. Responses to the light spot test were qualitatively similar across all three genotypes. On average, *Egr3*^-/-^ mice accumulated fewer wheel rotations, but this effect was not statistically significant (Fig. 4E).

### Heterozygous or Homozygous Deletions of Egr3 Ameliorate Ictal and Interictal Abnormalities in Kcna1^-/-^ mice

To directly test our hypothesis, we asked whether *Egr3* deletions can rescue epileptic and/or encephalopathic features in *Kcna1*^-/-^ mice. We bred double heterozygous *Egr3*^+/−^ *Kcna1*^+/−^ mice and studied the following four (out of 9 possible) genotypes: *Egr3*^+/+^ *Kcna1*^+/+^ (WT), *Egr3*^+/+^ *Kcna1*^-/-^, *Egr3*^+/−^ *Kcna1*^-/-^ and *Egr3*^-/-^ *Kcna1*^-/-^ mice. On this modified background, we observed that the loss of one or two copies of *Egr3* significantly prolonged survival (Fig. 5A). This effect was associated with an amelioration of overall hyperactivity levels (Fig. 5B, C). We did not observe reduced sheltering in *Egr3*^+/+^ *Kcna1*^-/-^ compared with *Egr3*^+/+^ *Kcna1*^+/+^ mice, and *Egr3*^-/-^ *Kcna1*^-/-^ displayed the lowest amount of sheltering overall (Fig. 5B, Fig. S3B). *Egr3* deletions also improved the duration of sucrose and water drinking bouts, without impacting overall lick numbers (Fig. 5D). Unlike hyperactivity patterns, behaviorally measured deficits in “sleep” were *not* restored by Egr3 deletion, although *Egr3*^-/-^ Kcna1^-/-^ mice did display more normalized “sleep” bout distributions (Fig. 5E). *Egr3* deletion did not alter responses to light spot stimulation (Fig. 5F), but did improve overall wheel running (Fig. 5G). Homozygous reductions in *Egr3* also facilitated significant reductions in overall GFAP immunoreactivity and hippocampal BDNF levels (Fig. 5H). To examine whether these neurobehavioral and histopathological changes were associated with improvements in seizure frequency, we conducted tethered EEG recordings that also permit measurements of spreading depolarization events in *Kcna1*^-/-^ mice.^125,126^ Over a 12-day recording period, heterozygous deletions of *Egr3* lowered the frequency of spontaneous seizures, and *Egr3*^-/-^ *Kcna1*^-/-^ mice were seizure free (Fig. 5I). Finally, to examine whether improvements in ictal and interictal behavioral features would be associated with a rescue in protein expression patterns, we conducted tandem mass tag proteomics on hippocampal samples from *Egr3*^+/+^ *Kcna1*^+/+^ (WT), *Egr3*^+/+^ *Kcna1*^-/-^, *Egr3*^+/−^ *Kcna1*^-/-^ and *Egr3*^-/-^ *Kcna1*^-/-^ mice. In a hierarchical clustering tree revealed from protein expression patterns, samples from *Egr3*^-/-^ *Kcna1*^+/+^ and *Egr3*^-/-^ *Kcna1*^-/-^ mice were most similar, and together, these genotypes most closely resembled expression patterns in WT (*Egr3*^+/+^ *Kcna1*^+/+^ mice).

**Figure 5.**
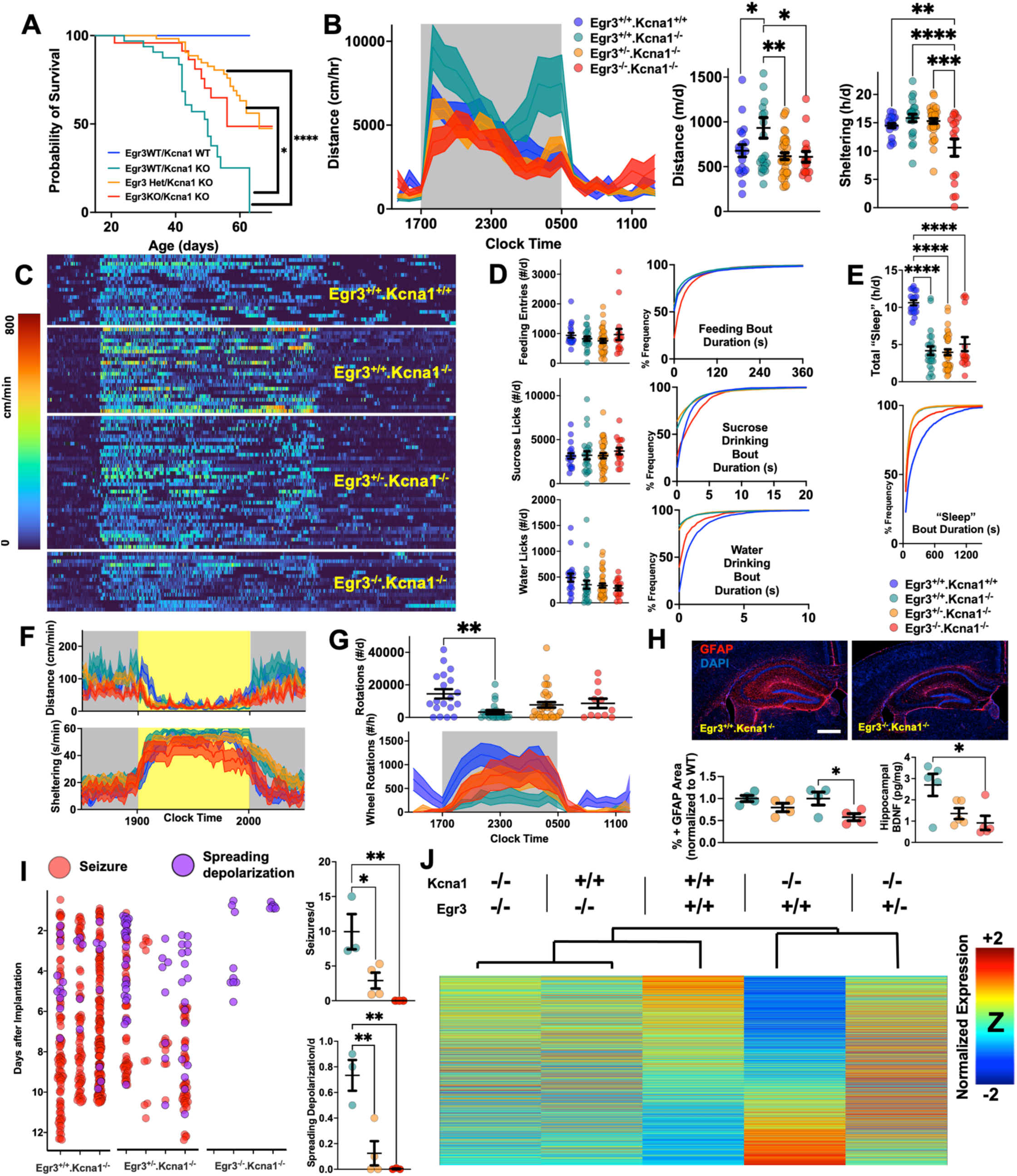
Egr3 Deletion Ameliorates the Ictal and Several Interictal Abnormalities in *Kcna1*-/- Mice. (A) Compared with *Egr3*+/+ *Kcna1*-/- mice (n=32), *Egr3*+/− *Kcna1*-/- mice (n=56) and *Egr3*-/- *Kcna1*-/- mice (n=24) displayed significantly prolonged survival. (B) As assessed through hourly distances, *Egr3*+/+ *Kcna1*-/- mice (n=23, 13 female) were significantly more hyperactive compared with WT (*Egr3*+/+ *Kcna1*+/+, n=19, 5 female), *Egr3*+/− *Kcna1*-/- (n=36, 19 female) or *Egr3*-/- *Kcna1*-/- mice (n=16, 6 female, F66, 1980 = 3.42, p<0.0001). Total distances for B (F3,90 = 4.44, p<0.001) and total sheltering (F3,90 = 8.36, p<0.0001). (C) Locomotor activity raster plot for all mice shown in B. (D) Total number of feeding, sucrose and water licking bouts for all four genotypes, together with cumulative frequency distribution histograms shown. Compared with *Egr3*+/+ *Kcna1*-/- (teal), *Egr3*-/- *Kcna1*-/- mice (red) displayed more prolonged bouts across all three categories (KS p<0.0001). (E) *Egr3* deletion did not improve reductions in total daily “sleep” in *Kcna1*-/- mice, but did partially improve “sleep” bout durations (KS p<0.0001). (F) Responses to light spot testing. (G) *Egr3* deletion rescued reductions in wheel-running in *Kcna1*-/- mice assessed in hourly bins (F69, 1734 = 1.31, p<0.01) or total wheel rotations (F3,79 = 4.52, p<0.001). (H) *Egr3* deletion also improved astrogliosis, as measured by GFAP immunoreactivity and total hippocampal BDNF levels. (I) Using conventional tethered equipped with DC band recording facilities, *Egr3*-/- *Kcna1*-/- mice and to a lesser extent, *Egr3*+/− *Kcna1*-/- mice featured a significant improvement in the frequency of spontaneous seizures and spreading depolarization events (n=3-4/group). (J) Hippocampal proteomic profiles from ∼7-week old mice visualized as heatmaps of relative Z-normalized expression (averaged *across* biological replicates), ranked in order of increasing fold change in *Egr3*-/- *Kcna1*-/- mice compared with *Egr3*+/+ *Kcna1*-/- mice. The dendrogram shown was constructed by hierarchical clustering of protein expression profiles using average linkage on correlation-based pairwise distances between biological samples. Mean + s.e.m shown for all. *, **, ***, **** depict p<0.05, <0.01, <0.001 and <0.0001 respectively. KS (Kolmogorov-Smirnov test).

## Discussion

In this study, we leveraged the advantages of home-cage monitoring to elaborately visualize a syndrome of neurobehavioral abnormalities that co-occur with spontaneous seizures in mice lacking *Kcna1*/Kv1.1.^39^ Under task-free conditions, *Kcna1*^-/-^ mice displayed marked hyperactivity and hyperarousal, together with excessively fragmented bouts of sleep, feeding and drinking. In response to light-spot stimulation within their own home-cages^26–29^, *Kcna1*^-/-^ mice displayed a relatively abrupt reduction in home-cage exploration, potentially reflecting enhanced risk aversion (paralleling states of heightened anxiety^66^). *Kcna1*^-/-^ mice also displayed fewer sucrose licks and accumulated fewer wheel rotations, pointing to alterations in natural reward sensitivity. Importantly, these quantitative phenotypes were acquired digitally through (i) prolonged recordings, (ii) conducted under experimenter-free conditions and (iii) *after* the habituation to a novel standardized cage that the mouse adopts as its home-cage.^20^ We propose that this constellation of changes in sleep/arousal, sensory responsivity, amotivation and behavioral fragmentation effectively models the *encephalopathy* of DEE.

In search for the upstream determinants of these encephalopathy-like behavioral changes, we sought to separate the neurobehavioral consequences of early widespread *Kcna1* loss from those purely downstream of spontaneous seizures (caused by the loss Kv1.1). Using wireless EEG deployed within home-cages, we found that daily measures of home-cage hyperactivity were *positively* correlated with daily variations in seizure frequency. Rather than infer any true causal relationship (i.e., seizures *cause* hyperactivity, or vice versa), this suggests that biological mechanisms which dynamically modulate seizure risk in Kcna1^-/-^ mice may pleiotropically alter peri-ictal home-cage behavior. To further isolate the contributions of seizures, we examined how variations in KCNA1 expression that do not result in spontaneous seizures impact home-cage behavior. Under identical home-cage conditions, mice that were haploinsufficient for *Kcna1* and which are seizure free^39^ were effectively asymptomatic. Mice with deletions of *Lgi1* (which express 50% less KCNA1 at axon initial segments^77^) only displayed mild insomnia without hyperactivity. In contrast, those with homozygous deletions of *Cntnap2* (which lack KCNA1 at juxtaparanodal regions^79^) featured *hypoactivity* without hypersomnia, and double heterozygous (*Lgi1*^+/−^ *Cntnap2*^+/−^) mice were also behaviorally asymptomatic. By utilizing *Kcna1^fl^*^/fl^ mice^43^, we found that tamoxifen-induced^64^ adult-onset deletions of *Kcna1* also only produced mild insomnia without impacting activity levels and bout structure. We further conducted a limited survey of cell-type specific *Kcna1* deletions: only PVCre+*Kcna1^fl^*^/fl^ mice displayed a mild behavioral phenocopy of *Kcna1*^-/-^ mice, featuring significant reductions in sheltering and total daily “sleep”, while deletions of *Kcna1* in Emx1-Cre or DAT-Cre expressing neurons did not alter home-cage metrics. Taken together, these results highlight how widespread KCNA1 loss occurring in early neurodevelopmental windows *and* early onset spontaneous seizures are *both* necessary to establish aberrations in home-cage behavior in *Kcna1*^-/-^ mice. These results provide a preclinical example of how the encephalopathy of this DEE has both lesional developmental *and* epileptic contributions. As a corollary, these data suggest that gene replacement strategies designed to alleviate DEE-like features in patients with KCNA1 DEE must be administered early in disease course *and* successfully transduce a wide population of cell types to be efficacious.

Next, we pivoted to exploring candidate molecular mechanisms *downstream* of Kcna1 loss *and* seizures that might perpetuate a state of pervasive seizure risk and encephalopathy. We chose to conduct a proteomic exploration of the hippocampus, a nidus for seizure generation in *Kcna1*^-/-^ mice, and where prominent astrogliosis^51^ and mossy fiber sprouting is also evident following status epilepticus.^40^ While several significantly regulated proteins were worthy of follow up, our attention was drawn to the induction of EGR3 and BDNF, given the extensive and highly corroborated studies demonstrating upregulations of EGR3 following electroconvulsive seizures. ^50,56,115,117,127^ *Increases* in *Egr3* mRNA levels (also measured cross-sectionally) have been observed following amygdala kindling^128^, and in three distinct rodent models of audiogenic seizures (the *tremor* mouse^129^, the Wistar Audiogenic Rat and the Genetic Audiogenic Seizure Hamster^130,131^). BDNF may function both upstream *and* downstream of EGR3: while Egr3 deletions severely attenuate BDNF inductions following electroconvulsive seizures ^56,117^, BDNF has been shown to induce EGR3 in ex vivo cortical neuron culture models.^132,133^ Using continuous video-EEG in rats rendered epileptic through pilocarpine injections, one previous study showed that mRNA levels for both BDNF and Egr3 were increased 3h following spontaneous seizures, and which both returned to baseline levels 24h later.^134^

To examine whether Egr3 deletion may ameliorate either neurobehavioral or epileptic features in *Kcna1*^-/-^ mice, we crossed double-heterozygous *Egr3*^+/−^*Kcna1*^+/−^ mice. Compared to WT mice in this modified background, (*Egr3*^+/+^)*Kcna1*^-/-^ mice still displayed spontaneous seizures, premature mortality, hyperactivity, insomnia, reduced wheel-running and fragmented sleep and licking bouts (Fig. 5), but did *not* display reduced sheltering, reduced sucrose-licking or abnormal light-spot responses (Fig. 1). Deleting one or two *Egr3* alleles led to improved seizure frequency, prolonged survival, attenuated hyperactivity and restored wheel-running interest in *Kcna1*^-/-^ mice but did not improve reductions in total daily “sleep”. Ablating Egr3 also improved markers of astrogliosis and reduced BDNF levels, suggesting that at least in the context of Kcna1 loss, BDNF induction may indeed be downstream of Egr3 induction. Multiple lines of evidence implicate a proconvulsant/pro-epileptogenic role for BDNF.^108^ Chemically induced status epilepticus dramatically raises BDNF levels and the activation of BDNF’s receptor, tropomysin-related kinase B (TrkB), whereas inhibiting TrkB post-status epilepticus reduces the frequency of spontaneous seizures and deficits in exploratory behavior.^135^ Mice with transgenic overexpressions of BDNF *or* TrkB display hippocampal CA3 hyperexcitability and experience more clinically severe kainate-induced seizures.^121,122^ Direct hippocampal infusions of BDNF have also been reported to elicit spontaneous limbic seizures.^136^ Finally, *Egr3* deletion also improved the frequency of spreading depolarization events which may correlate with improved survival in *Egr3*^-/-^ *Kcna1*^-/-^ compared with *Egr3*^+/+^ *Kcna1*^-/-^ mice.^137^ While transcription factors have been traditionally considered to be ‘undruggable’ targets^138^, inhibitors of Egr1 (whose zinc finger domains are 90% homologous to Egr3^139^) are currently being developed to treat atopic dermatitis, where Egr1 plays a crucial role in modulating local inflammation and cell replication.^140,141^ Since we observed significant disease modifying effects with deletion of only a single copy of *Egr3*, and since *Egr3*^+/−^ mice were phenotypically indistinguishable from WT mice under home- cage conditions, we expect that partial pharmacological inhibition of Egr3 that is instituted in early developmental windows could offer disease modifying effects that may adjunct routine antiseizure medications use.

We acknowledge four main limitations to this study. First, while our home-cage metrics within each experiment were consistently compared between littermate subjects, we did not back-cross our mice to ensure that all mice were studied on the same background. We transparently report these background effects: for example, Cre-negative *Kcna1*^fl/fl^ mice in our PV-Cre experiment were hyperactive compared with Cre-negative *Kcna1*^fl/fl^ mice within Emx1 comparisons. Similarly, WT littermates to *Lgi1*^+/−^ mice behaved distinctly when compared to WT littermates of *Kcna1*^-/-^ mice. This may have introduced potential floor or ceiling effects when examining U-shaped continuous variables such as distances/day, licks/day, etc. Second, we did not systematically conduct wired or tethered EEG recordings for most of our behavioral comparisons. We focused on behavioral metrics as a primary endpoint and therefore cannot rule out whether spontaneous seizures in any single mouse either disrupted or normalized our selected metrics. Third, while our home-cage approach provides a detailed analysis of the spatiotemporal structure of spontaneous behavior over a representative day, they still provide only a snapshot of behavior in singly housed mice. Solutions to achieve individual subject-level kinematic data from group housed mice over multi-day or multi-week recordings remains an active area of technological innovation. Fourth, while our home-cage survey provides quantitative metrics regarding several dimensions of mouse behavior (rest and activity patterns, risk aversion, natural reward, appetitive behaviors), it does not directly or indirectly measure aspects of learning and memory, which is itself a broad multidimensional set of constructs (fear learning, spatial learning, object learning, etc.). Nevertheless, home-cage behavioral measurements provide a valuable backdrop with which to appraise learning and memory deficits, if present. While memory impairments in Kcna1^-/-^ mice^142^ might not be *caused* by poor sleep or hyperactivity, pharmacological treatments to improve sleep function may improve memory function.

In summary, our results validate the use of home-cage monitoring as an ethologically sound approach to measuring neuropsychiatric comorbidity in a monogenic mouse model of DEE. We provide a set of precision endpoints that can be employed to test future therapies aimed at alleviating seizures and encephalopathy-like behavioral changes, and which can be compared across mouse models.^26,27,29^ By combining these endpoints with conventional EEG recordings, we show that targeted deletions of a seizure-induced transcription factor (EGR3) can achieve therapeutic improvements in both ictal and interictal features of disability.

## Data Availability

All raw data from this study are available from the corresponding author upon reasonable request. Mass spectrometry proteomics data has been deposited publicly (see supplemental methods).

## Supplemental Material

### Supplemental Methods

#### Mass Spectrometry Proteomics

Bilateral whole hippocampi were rapidly dissected from mouse brains between 1000-1400h, frozen in dry ice, stored at −80C and provided to the BCM Mass Spectrometry Proteomics Core Facility. We applied microscaled label-free proteome profiling to first assess proteomic disturbances between WT and *Kcna1*^-/-^ mice (n = 3/group, Fig. 4A/Supplemental Table 1), ^143^. Frozen tissues were pulverized on a cold metal block with mechanical action. Tissue powder was dissolved in 50mM ammonium bicarbonate and digested using LysC enzyme for 2 hours at room temperature followed by trypsin at 37^0^C overnight. Enzyme reaction was neutralized using 10% formic acid and peptides were measured using the Pierce™ Quantitative Colorimetric Peptide Assay (Thermo Scientific 23275). Offline high pH STAGE peptide fractionation (15 fractions combined to 5 peptide pools) for 50µg peptide was carried out as described previously^143^. LC-MS/MS analysis was carried out using a nano-LC 1000 system (Thermo Fisher Scientific, San Jose, CA) coupled to an Orbitrap Fusion mass spectrometer (Thermo Fisher Scientific, San Jose, CA). Peptides were eluted using a 110min gradient of 2-30% acetonitrile/0.1% formic acid at a flow rate of 200nl/min onto the mass spectrometer operating in data-dependent acquisition mode with 3 second cycle time and a 5 second dynamic exclusion. MS1 was acquired in Orbitrap (120,000 resolution, 350-1400m/z) followed by MS2 in IonTrap (HCD 30%, AGC 5E4, 30ms ion injection). Raw data were processed with Proteome Discoverer 2.1 software (Thermo Scientific) using Mascot 2.4 (Matrix Science^144^) against the mouse NCBI RefSeq database updated on 03/24/2020. Search settings included Trypsin/P enzymatic digestion to form peptides from 7-50 amino acids in length, a parent ion charge between 2-6, a maximum of 2 missed cleavages, and cleave protein n-term methionine enabled. Precursor and product ion tolerances were set to 20ppm and 0.5Da respectively. Dynamic modifications included oxidation on methionine, carbamidomethyl on cysteine, and protein N-terminal acetylation. Identified peptides were validated at 5% false discover rate (FDR) with Percolator^145^. The gene product inference and iBAQ-based quantitation was carried out using the gpGrouper algorithm^143^. Expression data was median-normalized before downstream data analyses. Differentially expressed proteins were calculated using a moderated t-test to calculate p-values and log_2_ fold change in the R package limma with multiple hypotheses correction by Benjamini-Hochberg method^146^.

To examine how proteomic changes may be restored by *Egr3* deletion (Fig. 5J/Supplemental Table 2), we simultaneously profiled identically collected hippocampal samples from *Egr3*^+/+^ *Kcna1*^+/+^ (n=5), *Egr3*^+/+^ *Kcna1*^-/-^ (n=4), *Egr3*^+/−^ *Kcna1*^-/-^ (n=3), *Egr3*^-/-^ *Kcna1*^-/-^ (n=4) and *Egr3*^-/-^ *Kcna1*^+/+^ mice (n=4). Given the more complex group design, tandem mass tag (TMT) labeling was chosen to reduce technical variability, as multiplexing allows for samples to be analyzed simultaneously, controlling for batch effects. Protein extraction, digestion and peptide fractionation was carried out as previously described^147^. Briefly, pulverized tissue was lysed in 8M urea and digested using LysC and Trypsin proteases as before, and 25µg peptide per sample were labeled with TMT10 plex isobaric label reagent (Thermo Fisher Scientific) according to the manufacturer’s protocol. High-pH offline fractionation was performed, combined into 24 peptide pools, and acidified to a final concentration of 0.1% formic acid. Deep-fractionated peptide samples were separated on a nano-LC 1200 system coupled to the Orbitrap Exploris480 mass spectrometer (Thermo Fisher Scientific). Peptides were eluted using a 110min gradient of 2-30% acetonitrile/0.1% formic acid at a flow rate of 200nl/min onto the mass spectrometer operating in data-dependent acquisition mode, with a 2 second cycle time and a dynamic exclusion of 20 seconds. MS1 was measured in the Orbitrap at 120000 resolution, scan range 375-1500m/z and 50ms Injection time, followed by MS2 in the Orbitrap at 30000 resolution (HCD 38%) with TurboTMT algorithm. Dynamic exclusion was set to 20 seconds and the isolation width was set to 0.7m/z. MS raw data processing and differential analysis was carried out as described previously^148^, with precursor AUCs and reporter ion intensities extracted with Masic^149^]. Reverse decoys and common contaminants were added to the GENCODE mouse protein database using Philosopher^150^. Mass spectra were searched using MSFragger (v3.5) with mass calibration^151^. Fixed modifications included carbamidomethylation of cystine (+57.021465), and TMT reagent on lysine (+229.1629). Variable modifications include oxidation of methionine (+15.9949), peptide N-term pyroGlu formation from Q (−17.02560) or E (−18.01060), peptide N-term 2-oxo-propanoic acid formation from C (−17.00329, approximated as – 17.02560), protein N-term acetylation (+42.01567), and peptide N-term TMT (+229.1629). Database search was performed with strict trypsin rule (cleave after K/R) for peptides between 7 and 50 amino acids, charge 2-6, between 350-10,000 Da, up to 2 missed cleaves, and cleave protein n-term methionine enabled. Precursor mass tolerance was set to 20 ppm, a fragment mass tolerance of 0.02 Da. The top 300 peaks were used, with a minimum of 15. Peptide validation was performed with Percolator and filtered to 1% PSM FDR^145^. Gene product inference and iBAQ-based quantification was carried out using the gpGrouper algorithm^143^. Expression data was median-normalized and batch corrected for TMT multiplex with ComBat^152^ before downstream data analyses. Differentially expressed proteins were calculated using the moderated t-test to calculate p-values and log fold changes as implemented in limma^146^.

#### Electroencephalography

Our study employed both wireless and wired/tethered EEG. To enable simultaneous home-cage monitoring and EEG recording and permit unhindered access to shelters (Fig. 1I, J), we implanted ∼5.5-week-old *Kcna1*^-/-^ mice with EMKA easyTEL S-ETA devices (*wireless*; under sterile precautions and isoflurane anesthesia), that included (i) two biopotential leads that will be affixed subdurally in right frontal and left posterior parietal regions using dental cement, tunneled to a (ii) a wireless transponder positioned in the left flank. Following a ∼96h recovery period (featuring meloxicam analgesia and saline boluses as needed), we transferred mice into home-cages with two wireless receivers placed underneath. At least 8 complete days of wireless EEG was acquired at 1000Hz through IOX2 (EMKA’s bio- potential signal acquisition interface) and reviewed using LabChart reader using a bandpass filter (1-30Hz). We tallied electrographic seizures, which were easily recognizable epochs of rhythmic EEG activity with evolutions in frequency and morphology. 96 days of combined home-cage wireless EEG recordings from *Kcna1*^-/-^ mice (n=15) were divided into seizure free or seizure prone days (containing at least one spontaneous seizure). To measure seizure occurrence in *Kcna1*^-/-^ mice with or without *Egr3* deletions, we employed a tethered approach positioning 36 gauge Teflon coated silver wires subdurally over mouse cortex and neck muscles, followed by a cement cap^125,126^. Following a five-day recovery period, EEG recordings were collected within observation cages furnished with bedding, *ad libitum* chow/water. EEG was recorded and analyzed using Bioamp DC amplifiers and LabChart. An aerial infrared camera permitted continuous video recording.

#### Immunohistochemistry (IHC) and Enzyme linked Immunosorbent Assay (ELISA)

IHC against EGR3 and GFAP was conducted using published protocols^153^ with slight modifications. Mice were deeply anaesthetized with an intraperitoneal injection of Beuthanasia-D [0.1ml of pentobarbital (390mg/L) and phenytoin (50mg/L)] and then transcardially perfused with cold phosphate buffered saline (PBS) followed by 4% paraformaldehyde (PFA)/ PBS. Brains were extracted and post-fixed in 4% PFA/PBS for 24h, and then dehydrated in 30% sucrose/PBS for 24h. After cryo-protection, brains were cautiously dipped in cryo-molds containing Tissue Plus optimal cutting temperature embedding medium (Fisher Healthcare) and frozen at −80°C. 20µm free-floating coronal cryosections were cut using a Leica CM1950 cryostat (Leica Biosystems) at −20°C. Prior to staining, sections mounted on Superfrost slides were allowed to warm to room temperature and washed with PBS [10 min], dipped in Liberate Antibody Binding solution (L.A.B. Polysciences Inc.) for antigen retrieval [5 min], followed by 3 x 5 min PBS washes, permeabilized with 0.5% TritonX- 100/PBS [30 min/37°C/water bath] and underwent an additional three 10 minute PBS washes. After drawing a PAP pen hydrophobic barrier, sections were blocked for an hour in 5% donkey serum/PBS and incubated in primary antibodies for 1 hour (37°C), followed by overnight room temperature incubations. Primary antibodies used were rabbit-anti-NeuN [Millipore #ABN78; 1:250; overnight/4°C], mouse-anti-GFAP (Clone N206A/8) [Neuromab #75-240; 1:500; overnight/4°C] and rabbit-anti-Egr3^115^; 1:1000; 48h/4°C]. Following this, sections were washed with PBS [10 min], incubated in Alexa Fluor 488 donkey anti-rabbit, Alexa Fluor 555 donkey anti-mouse and Alexa Fluor 568 donkey anti-mouse [Invitrogen; 1h*(except for anti-Egr3 which was 24h/4°C)*/RT/dark; 1:400]. Next, sections were thoroughly washed, incubated in DAPI [5 minutes, dark] and mounted with Prolong Diamond Antifade mountant [Invitrogen]. Coverslipped sections were imaged on a Keyence BZ-X800 fluorescent microscope. All images were captured at 20X and/or 4X magnification, and analyzed using FIJI software (ImageJ, NIH). For cell counting, images at 20X magnification were taken and cells were counted from 4-6 sections/mouse from a set of 4-5 mice/group. For ELISA measurements of BDNF protein levels, we employed an established commercially available Quantikine ELISA kit (R&D Systems) and followed the protocol as per manufacturer’s instructions^154,155^. Bilateral whole hippocampal samples were snap frozen and stored in −80°C. For preparing tissue lysates, hippocampi were thawed on ice, rinsed with 1X PBS, homogenized with equal volume of Lysis Buffer 17 (with Protease inhibitor cocktail [1:10]) and centrifuged at 13000g for 15 min at 4°C. Sample absorbance at 595nm wavelength was measured using microplate reader.

#### Quantitative Reverse Transcriptase Polymerase Chain Reaction

qRT-PCR measurements of *Kcna1* mRNA levels was performed as described previously^156^ with minor modifications. Dissected bilateral hippocampal specimens were snap frozen in dry ice and stored at −80°C. Tissues were thawed and homogenized in Trizol reagent (Sigma Aldrich, USA). Chloroform was added and centrifuged at 12,000 g/15min/4°C. Next, the aqueous layer was exposed to 100% 2- propanol and centrifuged at 12,000 g/10min/4°C. The pellet formed was washed in 75% ethanol twice, centrifuged 7500g/5min/4°C, dissolved in RNAse-free water and quantified through NanoDrop One (Thermo Scientific). Reverse transcription was conducted using a high-capacity cDNA-RT kit (Applied Biosystems), as per manufacturer protocols. Subsequently, quantitative real-time PCR (qRT-PCR) analysis was accomplished using Power SYBR Green PCR Master Mix (Applied Biosystems) on a Quant Studio 3 real-time PCR system (Applied Biosystems). We employed the ΔΔCT method, utilizing *Rplp0* as the reference transcript^157^.

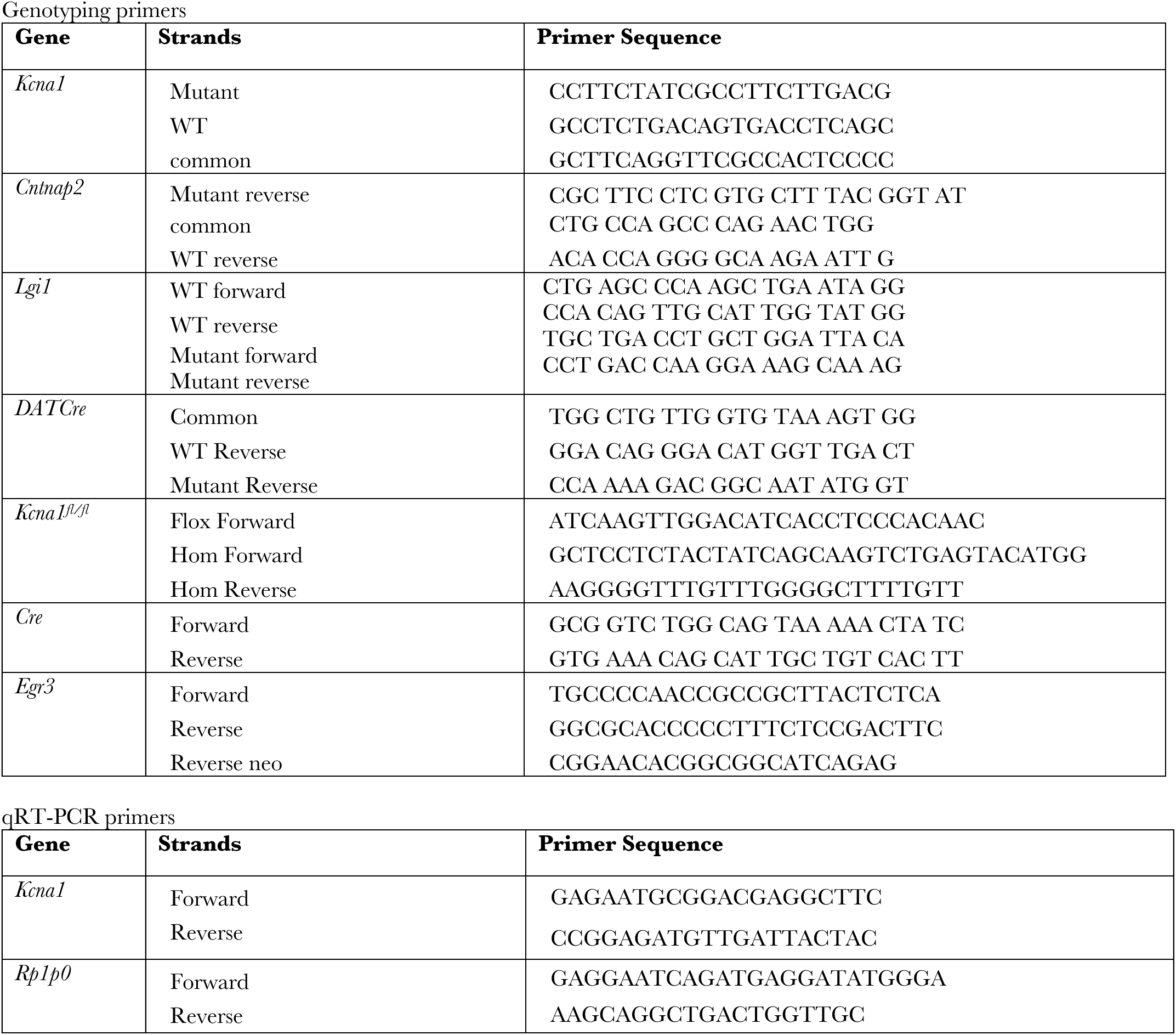

**Supplemental Figure 1.**
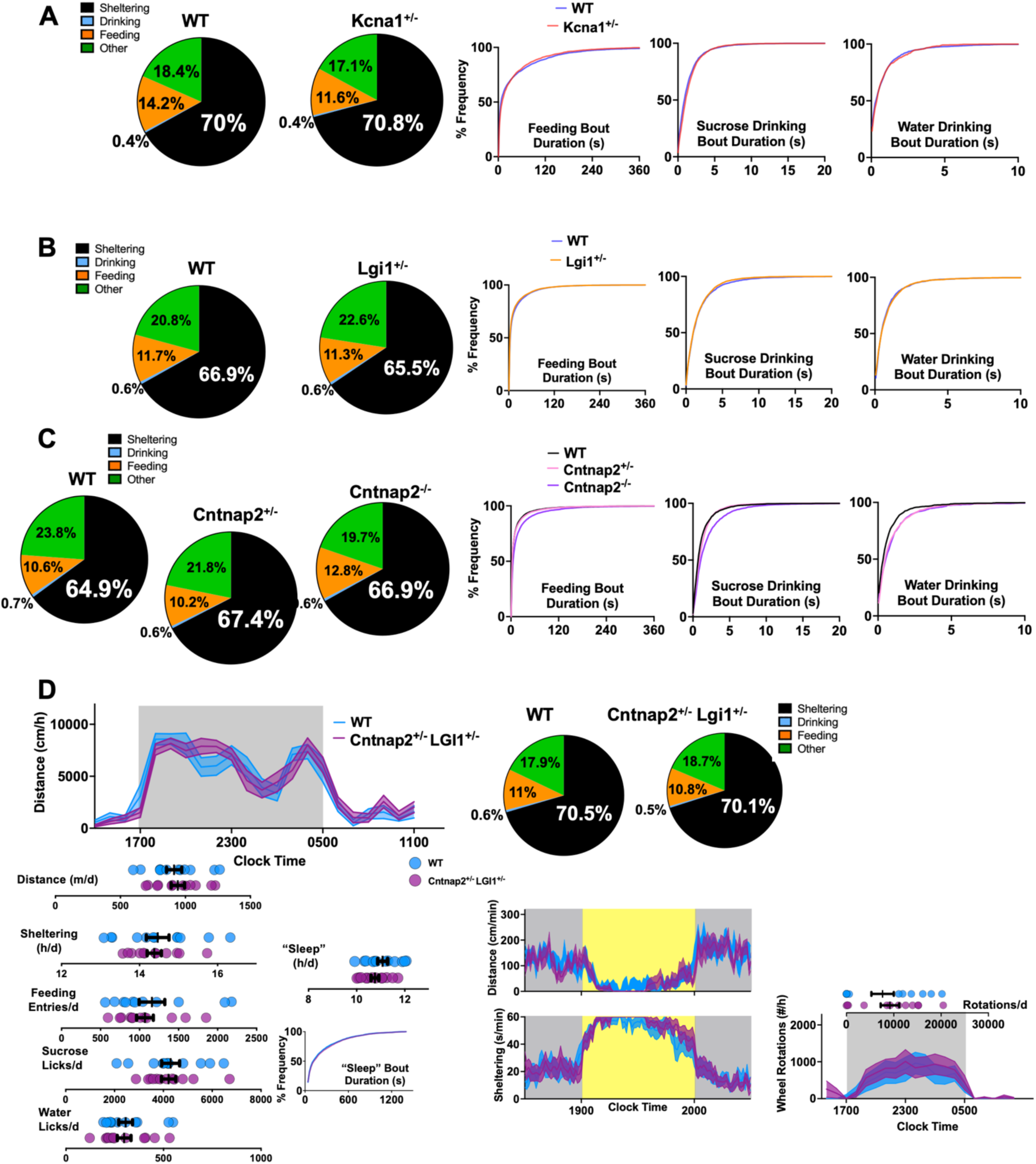
(A) Time budgets (of sheltering, feeding, drinking and “other” behaviors) and cumulative frequency distribution histograms of bout duration for WT and *Kcna1*+/− mice in Fig. 2A. (B) Time budgets and cumulative frequency distribution histograms of bout duration for WT and *Lgi1*+/− mice in Fig. 2B. (C) Time budgets and cumulative frequency distribution histograms of bout duration for WT, *Cntnap2*+/− and *Cntnap2*-/- mice in Fig. 2C. (D) *Cntnap2*+/− *Lgi*+/− mice double heterozygous mice behaved similarly to littermate WT mice under baseline conditions, light spot and wheel-running trials. Mean + s.e.m shown for all.

**Supplemental Figure 2:**
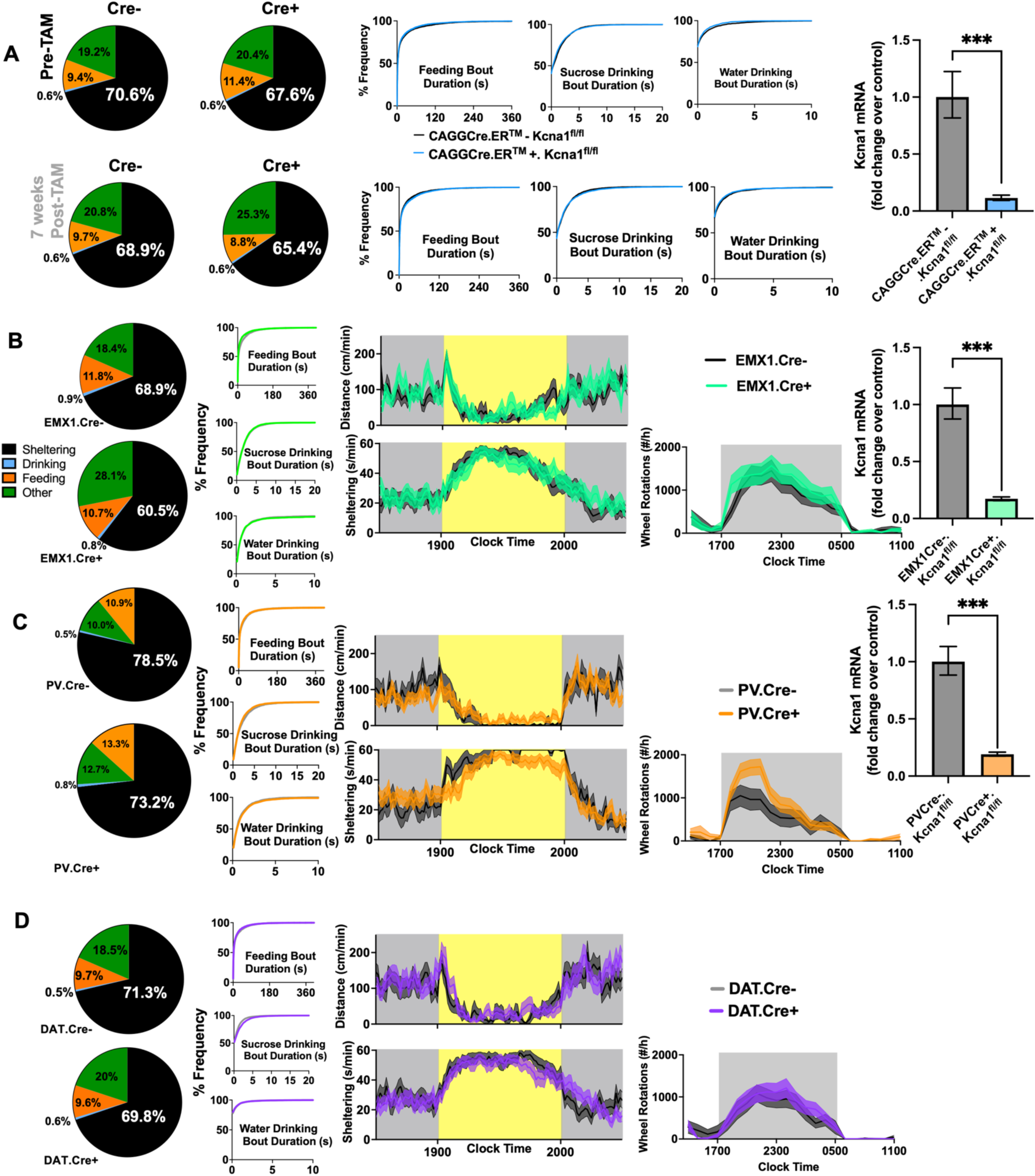
(A) Time budgets (of sheltering, drinking, feeding and “other” behaviors for CAGGCre.ERTM-positive and CAGGCre.ERTM-negative *Kcna1fl*/fl mice (Fig. 3A) before and 6 weeks after tamoxifen injections. Cumulative frequency distribution histograms of bout duration (center), and qRT-PCR validation of *Kcna1* mRNA knockdown (right). (B) Time budgets, cumulative frequency distribution histograms of bout duration, light spot, wheel-running trial and qRT-PCR validation of *Kcna1* mRNA knockdown in Emx1Cre-positive and Emx1Cre-negative *Kcna1fl*/fl mice (Fig. 3B). (C) Time budgets, cumulative frequency distribution histograms of bout duration, light spot, wheel-running trial and qRT-PCR validation of *Kcna1* mRNA knockdown in PVCre-positive and PVCre-negative *Kcna1fl*/fl mice (Fig. 3B). (D) Time budgets, cumulative frequency distribution histograms of bout duration, light spot and wheel-running trial in PVCre-positive and PVCre-negative *Kcna1fl*/fl mice (Fig. 3B). Mean + s.e.m shown for all. *** depicts p<0.001

**Supplemental Figure 3:**
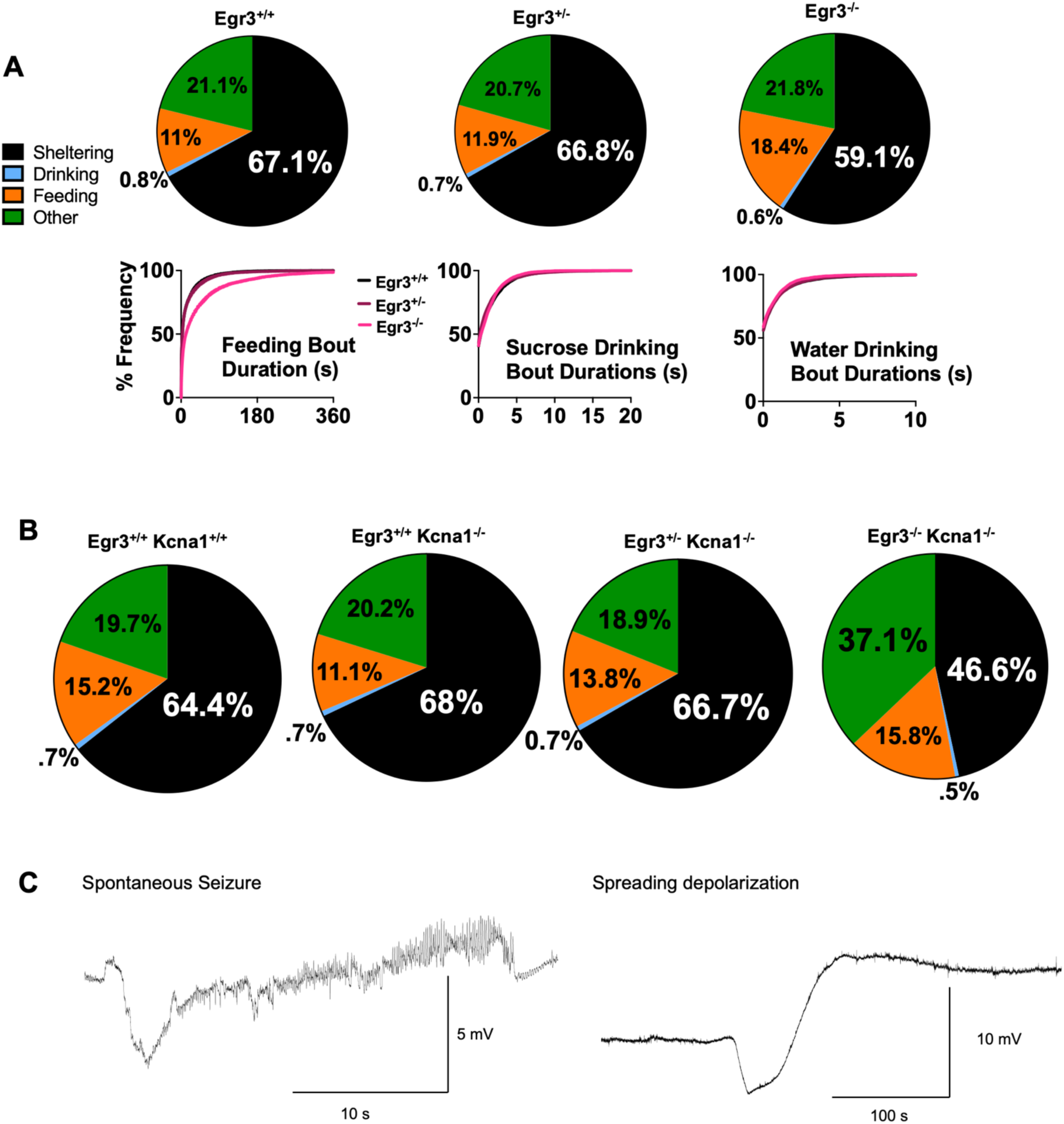
(A) Time budgets (of sheltering, drinking, feeding and “other” behaviors) and cumulative frequency distribution histograms of bout duration in *Egr3*+/+, *Egr3*+/− and *Egr3*-/- mice (Fig. 4E). *Egr3*-/- mice displayed more prolonged feeding bout durations (KS p<0.0001). (B) Time budgets for *Egr3*+/+ *Kcna1*-/- mice, *Egr3*+/+ *Kcna1*+/+, *Egr3*+/− *Kcna1*-/- and *Egr3*-/- *Kcna1*-/- mice. (C) Representative EEG traces (see Fig. 5I) capturing an electrographic seizure (left) versus an isolated spreading depolarization event (right). KS: Kolmogorov-Smirnov test.

